# The membrane activity of the amphibian Temporin B peptide analog TB_KKG6K sheds light on the mechanism that kills *Candida albicans*

**DOI:** 10.1101/2022.06.15.496139

**Authors:** Anant Kakar, Luis Enrique Sastré-Velásquez, Michael Hess, László Galgóczy, Csaba Papp, Jeanett Holzknecht, Alessandra Romanelli, Györgyi Váradi, Nermina Malanovic, Florentine Marx

## Abstract

Temporin B (TB) is a 13 amino acid long, cationic peptide secreted by the granular glands of the European frog *Rana temporaria*. We could recently show that the modified TB peptide analog TB_KKG6K rapidly killed planktonic and sessile *Candida albicans* at low µM concentrations and was neither hemolytic nor cytotoxic to mammalian cells *in vitro*. The present study aimed to shed light into its mechanism of action, with a focus on its fungal cell membrane activity. We utilized different fluorescent dyes to prove that it rapidly induces membrane depolarization and permeabilization. Studies on model membrane systems revealed that the TB analog undergoes hydrophobic and electrostatic membrane interactions showing a preference for anionic lipids and identified phosphatidylinositol and cardiolipin as possible peptide targets. Fluorescence microscopy using FITC-labelled TB_KKG6K in the presence of the lipophilic dye FM4-64 indicated that the peptide compromises membrane integrity and rapidly enters *C. albicans* cells in an energy independent manner. Peptide treated cells analyzed by cryo-based electron microscopy exhibited no signs of cell lysis; however, subcellular structures were disintegrated, suggesting that intracellular activity may form part of the killing mechanism of the peptide. Taken together, this study proved that the TB_KKG6K compromises *C. albicans* membrane function, which explains the previously observed rapid, fungicidal mode of action and promises its great potential as a future anti-*Candida* therapeutic.

**Importance:** Fungal infections with the opportunistic human pathogen *C. albicans* are associated with high moratility rates in immunocompromised patients. This is partly due to the yeast’s ability to rapidly develop resistance towards currently available antifungals. Small, cationic, membrane-active peptides are promising compounds to fight against resistance development as many of them effectuate rapid fungal cell death. This fast killing is believed to hamper the development of resistance, as the fungi do not have sufficient time to adapt to the antifungal compound. We prevously reported that the synthetic variant of the amphibian Temporin B peptide, TB_KKG6K, rapidly kills *C. albicans*. In the current study, the mechanism of action of the TB analog was investigated. We show that this TB analog is membrane-active and impairs cell membrane function, highlighting its potential to be developed as an attractive alternative anti-*C. albicans* therapeutic, which may hinder the development of resistance.

## Introduction

Fungal infections (mycoses) range from superficial infections of, e.g. the skin, nails and mucous membranes, to systemic infections that enmesh a number of organs such as the brain, heart, lungs, liver, spleen, and kidneys. Whilst the former category of fungal infections are not fatal and relatively straightforward to treat, the latter could be life threatening, especially in the case of immunocompromised patients, whose number is increasing due to heightened immunosuppressant usage, necessitated by the magnitude of transplant recipients and patients undergoing chemotherapy (1, 2). Currently 150 million severe cases of mycoses occur worldwide each year, resulting in around 1.7 million deaths (3). The opportunistic human pathogenic yeast *Candida albicans* is responsible for about 70% of fungal infections globally, making it the most common causative agent of mycoses in humans. Alarmingly, it is associated with mortality rates of over 40%, even with antifungal intervention (4).

The currently available antifungal drug repertoire is limited, and this dismal scenario is aggravated by the rise of resistance in fungi towards the major drug classes used as first line therapy (5).

Antimicrobial peptides (AMPs) are currently the subject of extensive investigation as a promising alternative therapeutic modality against microbial infections. AMPs, the majority of which are small, cationic peptides that are essential components of the innate immune response of many organisms, are effective against a variety of human pathogenic fungi (5, 6).

Temporins are secreted by the granular glands of the European red frog *Rana temporaria* and form one of the largest families of amphibian AMPs. They are short (8-14 amino acids), mildly cationic (0 to +3 at pH 7) and hydrophobic (∼50% hydrophobicity) peptides which are predominantly active against Gram-positive bacteria (7, 8). Recently, several modifications were made to the primary structure of the membrane-active peptide Temporin B (TB) in order to increase its efficacy and broaden its antimicrobial spectrum (9, 10, 11). The peptide analog that resulted in the highest growth inhibitory activity against both Gram-positive and Gram-negative bacteria was TB_KKG6K (TB analog; amino acid sequence: KKLLPIVKNLLKSLL; MW 1,718.2 Da); this contains four lysine residues, of which two were added to the N-terminus and one replaced the glycine (position 6) of the parent peptide. The new TB analog exhibits an increased positive net charge (from +0.9 to +3.9 at pH 7), reduced hydrophobicity (from 61% to 53%) as well as a modified amphipathic profile (10). Previous studies on synthetic TB_KKG6K were primarily focused on its antibacterial activity (10); however, its antifungal potential was less thoroughly investigated. We therefore evaluated its efficacy against the opportunistic human pathogen *C. albicans* only recently (7). We demonstrated that the peptide acted in a rapid and fungicidal manner against planktonic and sessile *C. albicans* cells, facilitating efficacious cell killing at low µM concentrations. In addition, we were able to demonstrate that this peptide is well tolerated by primary human skin cells and 3D skin epidermal models *in vitro*, indicating its potential to be developed as a therapeutic alternative against *C. albicans* skin infections (7).

The TB analog’s established antifungal efficacy encouraged us to further investigate this peptide in the current study by shifting our attention to its mechanism of action. We chose to focus on its activity on the fungal cell membrane because of its small size, positive charge, and rapid, fungicidal mode of action (7), which are characteristic features of cell membrane-active AMPs (12, 13). A multidisciplinary approach revealed that the TB analog compromised membrane function and rapidly entered the *C. albicans* cells where it disintegrated subcellular structures. The obtained results prove that this peptide is a promising, membrane-active amphibian biomolecule with the potential to join the next generation of peptide anti-*Candida* therapeutics.

## Results

### Determination of the growth inhibitory concentration of the test compounds

The inhibitory concentration that reduces the growth of *C. albicans* (CBS 5982) ≥ 90% (IC_90_) was determined for TB_KKG6K, octenidine (positive control), and PCγ^C-terminal^ (negative control) in broth microdilution assays using two yeast cell concentrations (1×10^4^ and 1×10^6^ cells/mL), as some experiments required different cell numbers. TB_KKG6K had an IC_90_ of 2 µM irrespective of the cell concentration used. This value corresponded to the previously reported IC_90_ determined against *C. albicans* by our group (7). Octenidine yielded an IC_90_ of 1 µM with 1×10^4^ cells/mL and 2 µM with 1×10^6^ cells/mL. As described previously (14), the peptide PCγ^C-terminal^ (amino acid sequence: CGGASCRG; MW 709.8 Da), which derives from the C-terminal part of the *Penicillium chrysogenum* antifungal protein C (PAFC), was found to be inactive against *C. albicans.* No fungal growth inhibition at the administered peptide concentration (0-32 µM) was detected.

### TB_KKG6K impairs the cell membrane integrity of *C. albicans*

To investigate how TB_KKG6K affects the integrity of the cell membrane of *C. albicans*, we first tested if the TB analog disrupts membrane polarity. To this end, we used the voltage-sensitive fluorescent 3,3′-dipropylthiadicarbocyanine iodide (DiSC_3_(5)) dye, which dimerizes and accumulates in the intact polarized cell membrane of energized cells, resulting in quenching of the fluorescent signal. Upon membrane depolarization, the DiSC_3_(5) dimers dissociate and are released into the supernatant. The release can be measured as an increase in fluorescence intensity (15). The membrane perturbing surfactant Triton X-100 was used as a positive control at a concentration of 1% (w/v). It induced a fast and sustained increase in fluorescence intensity, proving that it dissipated the cell membrane potential in a timely manner (Figure 1). PCγ^C-terminal^ was applied at 32 µM and served as the negative control. No membrane depolarization was observed with PCγ^C-terminal^ (Figure 1). The TB analog was applied in three different concentrations, 0.5 µM, 1 µM, and 2 µM. TB_KKG6K induced membrane depolarization in a concentration and time-dependent manner. The fluorescence intensity of the sample immediately increased upon the addition of 2 µM peptide (corresponding to the IC_90_ value) within the first 25 min of exposure. The response was less pronounced when the peptide was added at the sub-inhibitory concentration of 1 µM, and minimal with 0.5 µM (Figure 1). This result indicates that TB_KKG6K dissipates the cell membrane potential, which might be one reason for its growth inhibitory activity against *C. albicans*.

**Figure 1.**
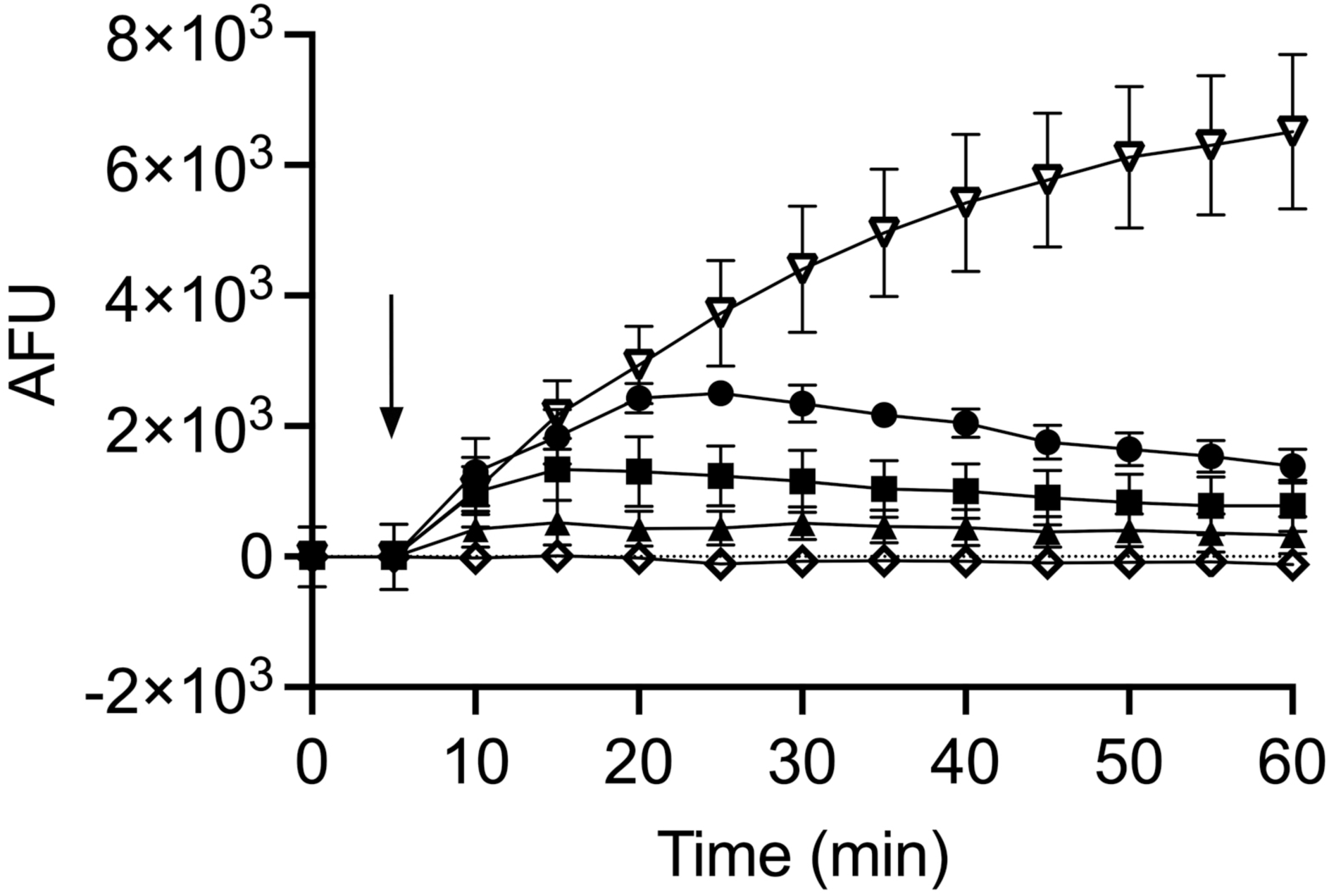
Membrane depolarization potential of TB_KKG6K in *C. albicans* determined with the DiSC_3_(5) dye. TB peptide, 0.5 µM (▲), 1 µM (▪), 2 µM (●); Triton X-100, 1% (v/v) (▽); PCγ^C-terminal^, 32 µM (◇). Compound addition is marked with an arrow and the depolarization of the cell membrane was monitored over a 60 min time period. The arbitrary fluorescence units (AFU) shown have been normalized by subtracting the background fluorescence values of the *C. albicans*-DiSC_3_(5) combination without compound addition (untreated control). The values represent the mean ± standard deviation (SD) of fluorescence values collected from two independent experiments performed in technical duplicates.

Next, we questioned, if the depolarization of the cell membrane by the TB analog coincides with its permeabilization. We, therefore, tested the membrane permeabilizing capacity of TB_KKG6K with the SYTOX Green uptake assay. The fluorescent SYTOX Green dye is a membrane impermeable molecule that penetrates only compromised cell membranes and fluoresces upon binding to nucleic acids (16). This was confirmed by the use of the positive control octenidine, which is a membrane-active compound previously shown to possess anti-*Candida* activity (17). Octenidine increased the fluorescence intensity of the sample after its addition at its IC_90_ value (2 µM) (Figure 2). TB_KKG6K was applied at 0.5 µM, 1 µM, and 2 µM and the change in fluorescence intensity after peptide addition was monitored for 60 min (Figure 2). Within the first 10 min of incubation, the rise in fluorescence intensity induced with 2 µM peptide was the fastest, followed by that of the sample exposed to 1 µM peptide. Steady-state signal intensity values were reached after 15-20 min of incubation with both peptide concentrations, which settled thereafter at a level similar to that of the positive control octenidine. The permeabilization of the membrane was delayed and less effective when 0.5 µM peptide was used. The negative control PCγ^C-terminal^ showed no membrane permeabilizing activity (Figure 2). Of note, the negative fluorescence values recorded with PCγ^C-terminal^ could be explained by the peptides’ possible interference with SYTOX Green, thereby preventing the interaction of the dye with the cell. The fact that the exposure of cells to TB_KKG6K resulted in a fast depolarization and permeabilization of the cell membrane indicates that the TB analog quickly kills *C. albicans* by compromising the cell membrane integrity.

**Figure 2.**
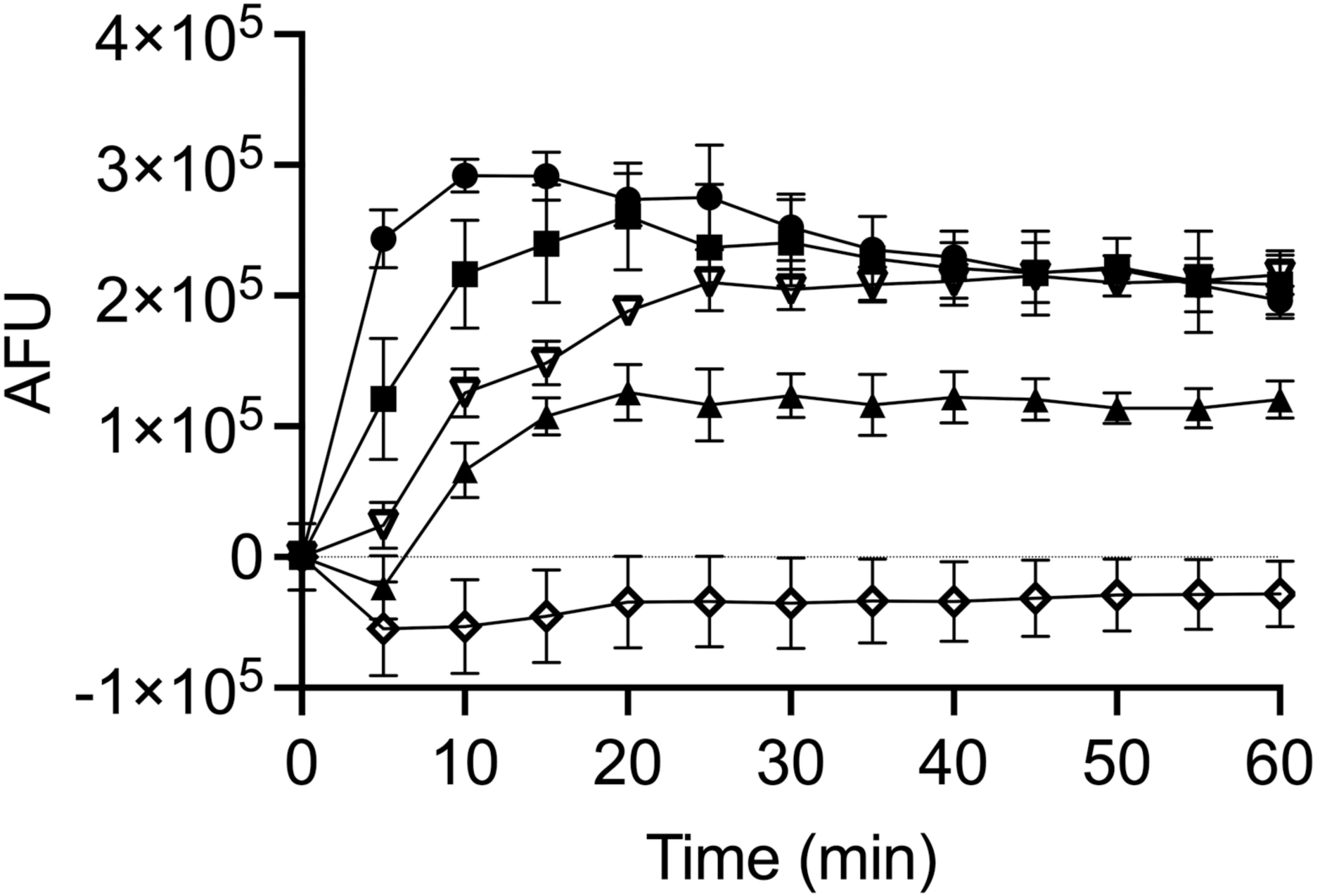
Membrane permeabilization activity of TB_KKG6K in *C. albicans* detected with the SYTOX Green uptake assay. TB peptide, 0.5 µM (▴) 1 µM (▪), 2 µM (●); octenidine, 2 µM (▽); PCγ^C-terminal^, 32 µM (◇). AFU were normalized by subtracting the background fluorescence of the medium with/without compounds in the absence of cells. AFU values depicted in the graph were further corrected by subtracting the fluorescence values of the *C. albicans*-SYTOX Green combination without compound addition (untreated control). The values presented are the mean ± SD determined from two independent experiments performed in technical duplicates.

### TB_KKG6K binds to phosphoinositide phosphates and preferentially permeabilizes anionic model membranes

Many cationic effector peptides of natural and synthetic origin were reported to be specifically attracted by negatively charged membrane components e.g. phospholipids, and these interactions have been shown to regulate their mode of antimicrobial action (18). We therefore tested the TB analog for its ability to bind phospholipids and performed a peptide-lipid overlay experiment using commercially available phosphatidylinositide phosphates (PIP) strips. Since antibodies directed against TB_KKG6K were not available, we used the peptide labeled with the green fluorophore fluorescein isothiocyanate (FITC; FITC-TB_KKG6K) and recorded the fluorescence intensity of the signal of the lipid-bound peptide on the membrane. As shown in Figure 3A, the peptide exhibited a preferential binding to lipids with higher negative charge, namely phosphatidylinositol (PI) mono-, bi- and tri-phosphates. Semi-quantification of the spot signal intensity on the membranes (n=3) confirmed the preference of TB_KKG6K for phosphorylated PI, but also for unphosphorylated PI (Figure 3B).The binding to phosphatidic acid (PA) and phosphatidylserine (PS), however, was not significantly higher compared to the blank (Figure 3B). No binding of the peptide occurred with lysophosphatidic acid (LPA), lysophosphocholine (LPC), phosphatidylethanolamine (PE), phosphatidylcholine (PC), or sphingosine-1-phosphate (S1P) (Figure 3A, B).

**Figure 3.**
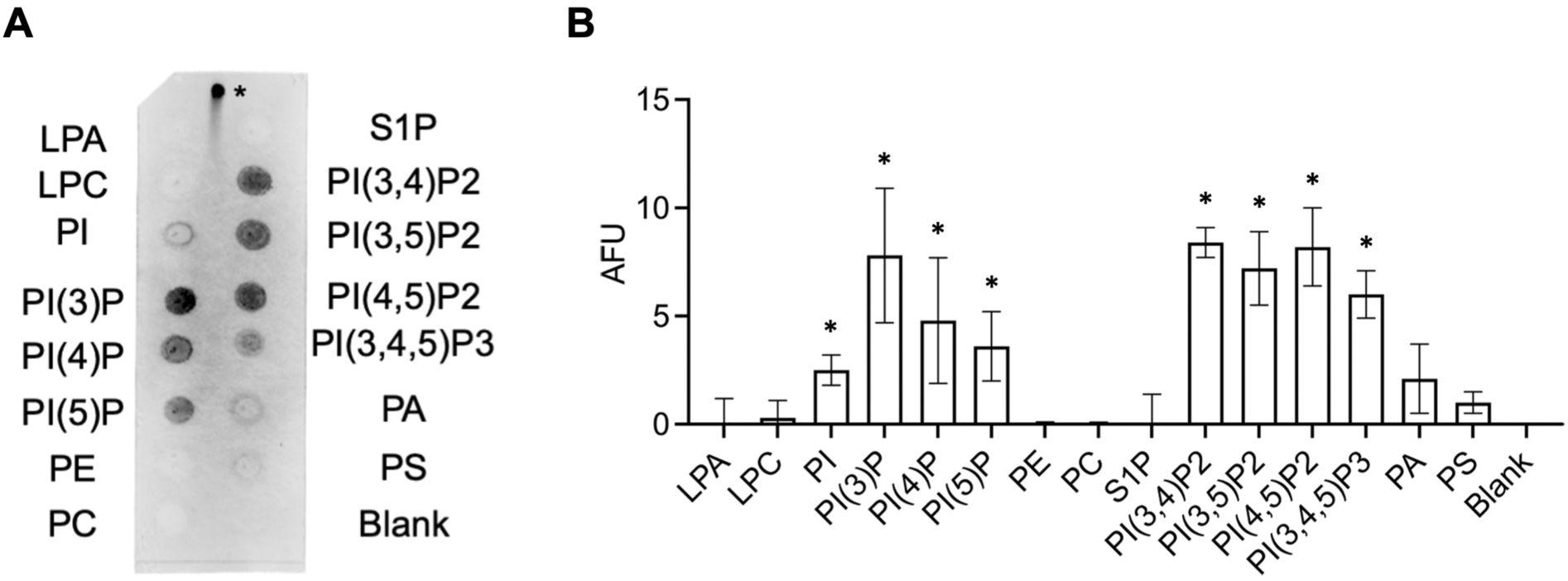
Binding of FITC-TB_KKG6K to phosphoinositide phosphates. (A) The PIP strip was probed with 1.5 µg/mL of FITC-labelled TB_KKG6K and the binding of the peptide to the lipids was fluorometrically detected. The positive control for fluorescence signal detection is marked with an asterisk and represents 0.5 µg of peptide spotted onto the membrane. The PIP strip shown represents the result of one out of three independent experiments. (B) Relative quantification of signal intensities of the spots representing TB_KKG6K bound to lipids compared to the blank, which was set as 0 arbitrary fluorescence units (AFU). AFU represent the mean ± SD of fluorescence values quantified in three independent blots by ImageJ/FIJI; \**p* ≤ 0.05. LPA, lysophosphatidic acid; LPC, lysophosphocholine; PIP, phosphoinositide phosphates; PE, phosphatidylethanolamine; PC, phosphatidylcholine; S1P, sphingosine-1-phosphate; PA, phosphatidic acid; PS, phosphatidylserine.

To gain a more detailed insight into the binding preferences of TB_KKG6K for certain lipids that influence its interaction with the cell membrane, we evaluated the peptide’s permeabilizing potential by using large unilamellar vesicles (LUVs) of different lipid composition and charge that represented prokaryotic and eukaryotic model membranes. The LUVs were loaded with 8-aminonaphthalene-1,3,6-trisulfonic acid and p-xylene-bis-pyridinium bromide (ANTS-DPX), which, upon the application of a permeabilizing compound, leaks out into the surrounding buffer and can be quantified fluorometrically (19–21). These changes in fluorescence intensity were followed in real-time before and 30 min after the addition of the compounds to be examined. The negative and positive controls used in this experiment were 32 µM PCγ^C-terminal^ and the membrane-lysing surfactant agent Triton X-100 (1% (w/v)), respectively. The fluorescence values that were reached due to the leakage of ANTS-DPX of LUVs exposed to the positive control represented 100% leakage. No membrane activity (≤ 2% leakage) was induced in LUVs by the negative control PCγ^C-terminal^ (Table 1). The peptide was applied in three concentrations, 2 µM, 4 µM, and 8 µM (representing a lipid-to-peptide molar ratio of 25:1, 12:1 and 6:1), respectively (Table 1).

**Table 1.**
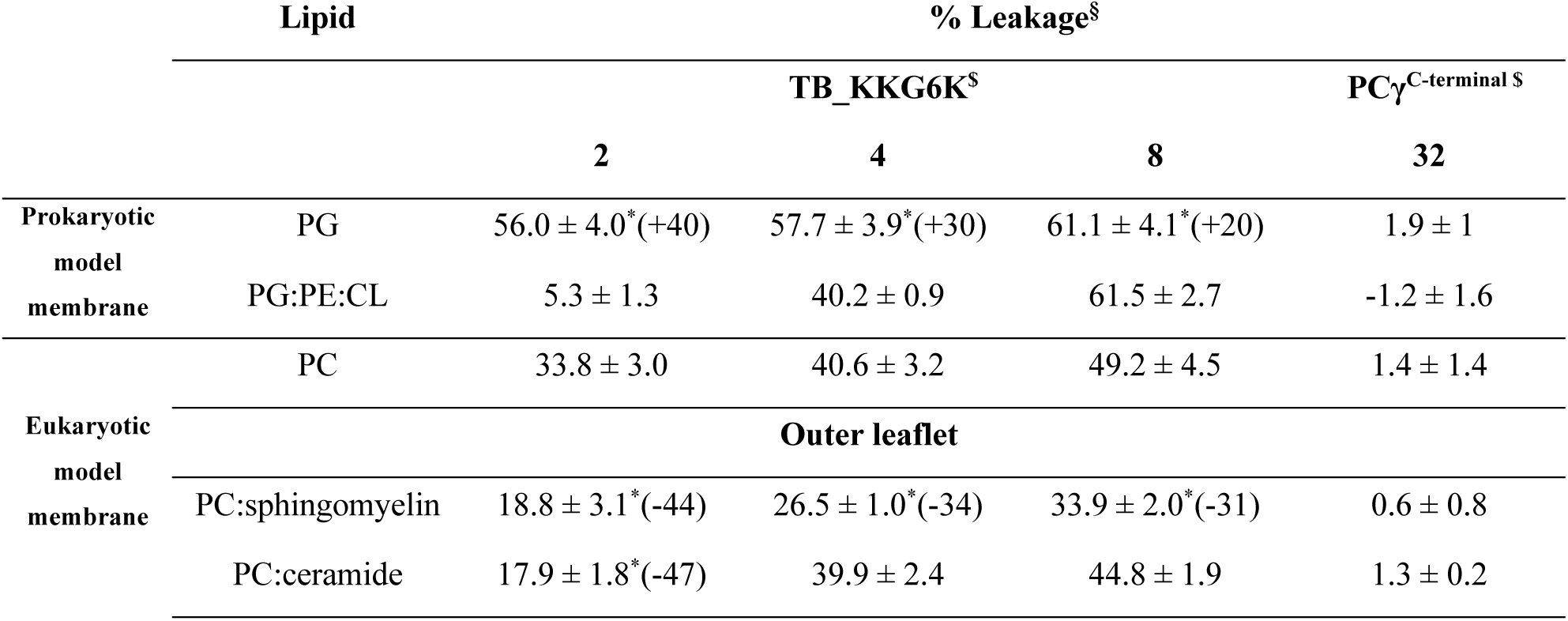
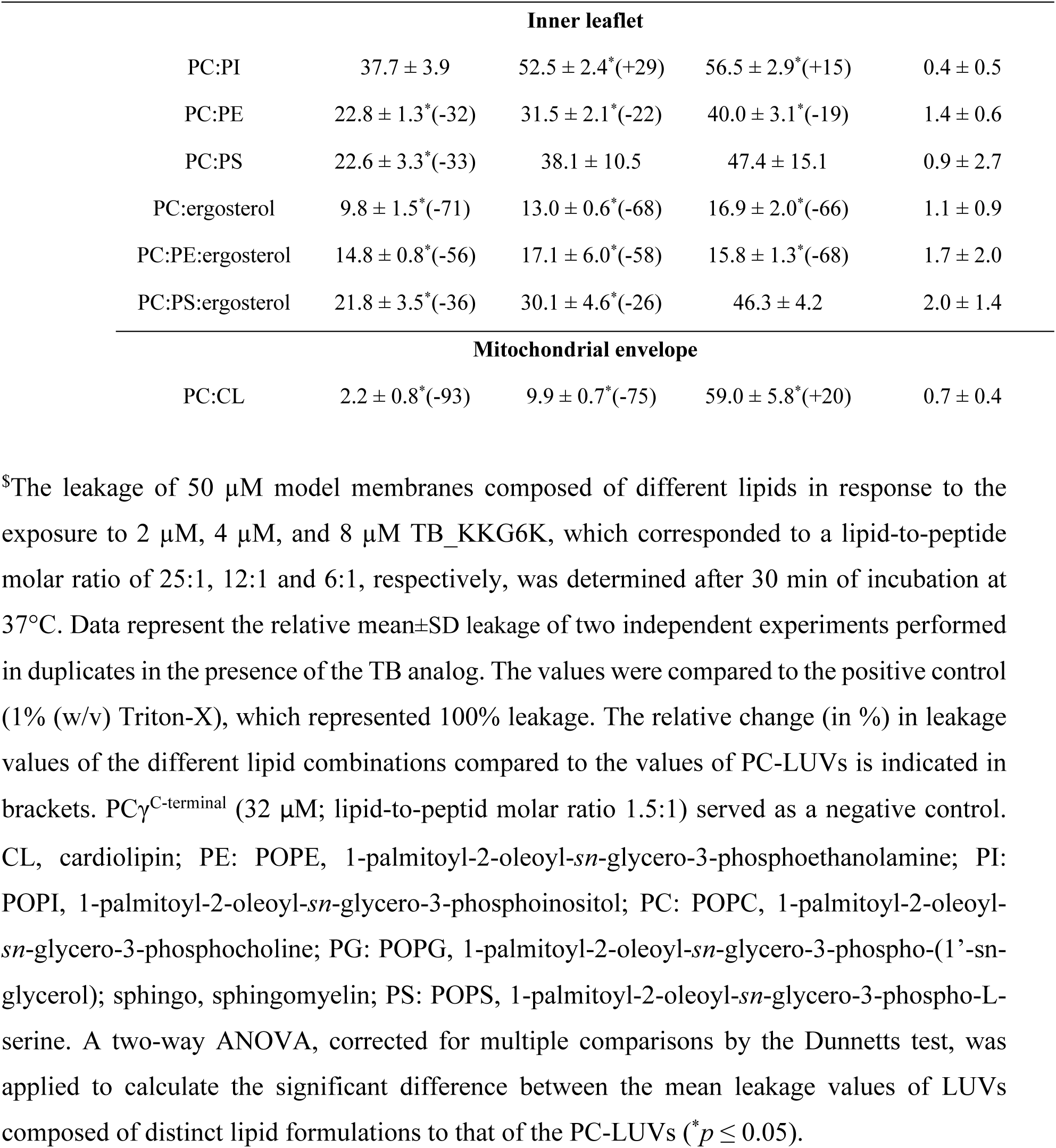
TB_KKG6K induced leakage of microbial model membranes.

Due to the well-characterized cell membrane activity of TB on bacteria, we first performed a control experiment and tested two model membranes that were composed of the main constituents of bacterial membranes, the anionic phosphatidylglycerol (PG), cardiolipin (CL) and the zwitterionic PE (22–25). The LUVs contained either PG alone (100% anionic), mimicking the membrane of Gram-positive bacteria, or a mixture of PG, PE and CL (PE:PG:CL, 30% anionic), representing the membrane of Gram-negative bacteria. As expected, TB_KKG6K interacted with both model membranes (Table 1). The leakage of LUVs composed of the less anionic lipids PE:PG:CL was strongly concentration dependent, whereby 8 µM peptide was required to reach 61±3% membrane leakage. In contrast, model membranes consisting of only PG were affected at a much lower peptide concentration; the exposure to 2 µM peptide resulted in 56±4% leakage, a value that increased only slightly with 8 µM peptide (61±4 %). This indicated that the maximum possible peptide-induced leakage of LUVs was reached in this experimental setting and pointed towards the peptide’s preference for anionic membranes (Table 1).

Next, we wanted to evaluate whether the TB analog prefers anionic membranes over neutral ones. Therefore, we compared the leakage elicited in LUVs composed of anionic PG to those containing the neutral phosphatidylcholine (PC). As shown in Table 1, TB_KKG6K interacted with PC and induced leakage in a concentration-dependent manner; however, the leakage values induced at all peptide concentrations tested were significantly higher (\**p* ≤ 0.05) with PG than with PC (difference in leakage with PG compared with PC: +40% at 2 µM; +30% at 4 µM; +20% at 8 µM). Although the peptide favored interaction with the anionic PG, the fact that it induced leakage in the zwitterionic PC as well suggests that not only electrostatic but also other forces, such as hydrophobic or hydrogen bonding, are involved in the interaction with the cell membrane lipids.

Finally, we tackled the question if the TB peptide preferentially targets any specific membrane lipid. To this end, we performed leakage studies using nine eukaryotic model membranes composed of PC along with one or two more lipids specific to eukaryotic membranes, namely CL, ceramide, ergosterol, PE, PI, PS, and sphingomyelin. PC formed the dominant part of these LUVs, as this neutral phospholipid is a major component of fungal and mammalian cell membranes (26, 27).

The leakage measurements revealed that among all the tested eukaryotic membrane formulations, the sensitivity of LUVs towards 2 µM and 4 µM TB_KKG6K was higher than that with PC alone only when this neutral lipid was combined with PI (increase in leakage compared to PC alone: +29% at 4 µM; +15% at 8 µM). The leakage of LUVs containing PC:CL was also higher (+20%) than that of PC alone in the presence of 8 µM peptide. Membranes composed of PC and ceramide, PE or PS did not yield higher leakage values when compared to that observed with PC-LUVs (Table 1). This let assume that these lipids do not represent specific targets of TB_KKG6K.

Of note, the presence of ergosterol or sphingomyelin in PC hampered the membrane activity of TB_KKG6K and inhibited the leakage of ANTS-DPX from LUVs composed of PC:sphingomyelin and PC:ergosterol, respectively (Table 1). The addition of ergosterol to zwitterionic PC:PE or charged PC:PS similarly inhibited the leakage of PC:POPE:ergosterol or PC:PS:ergosterol, but not of PC:PS:ergosterol when exposed to the highest peptide concentration applied (8 µM), suggesting that the anionic phospholipid PS can compensate for ergosterol’s inhibitory effect on leakage induction (Table 1).

### The physicochemical properties of TB_KKG6K favor membrane partitioning

The Membrane Protein Explorer software, mPEX v.3.3.0, was applied to investigate the membrane partitioning properties of the TB_KKG6K *in silico*, using the antimicrobial peptide LL-37 (residues 13-37 were applied to compute parameters, LL-37_13-37_) as a reference. This well-characterized AMP has previously been shown to be membrane-active and toxic to *C. albicans* (28). Figure 4 depicts the helical wheel projections of both peptides, indicating that both possess localized hydrophobic regions that could facilitate their interaction with and insertion into the hydrophobic membrane bilayer. This hydrophobic region, however, is larger in the TB analog (53% hydrophobicity) as it consists of eight uncharged, hydrophobic residues, in comparison to the five hydrophobic amino acids in LL-37_13-37_ (40% hydrophobicity), indicating that the TB_KKG6K might be able to insert deeper into the membrane compared to LL-37_13-37_. The TB analog, however, contains only four charged residues in its predicted α-helix, compared to six in LL-37_13-37_, implying that TB_KKG6K could undergo electrostatic interactions with the negatively charged membrane, but that the membrane attraction is less pronounced than in the case of LL-37_13-37_. According to the bilayer partitioning free energy calculation, the TB analog requires less energy to enter the bilayer (3.6 kcal/mol) compared to LL-37_13-37_ (20 kcal/mol) (Figure 4). Calculation of the amphipathicity of the respective peptide α-helices revealed that the hydrophobic moment of the TB analog was lower (13 µ) than that of LL-37 (21 µ) (Figure 4). This implies that the hydrophobic and hydrophilic residues are more evenly distributed among the helix of TB_KKG6K, resulting in increased amphipathicity, which in turn, facilitates membrane binding and insertion (29, 30).

**Figure 4.**
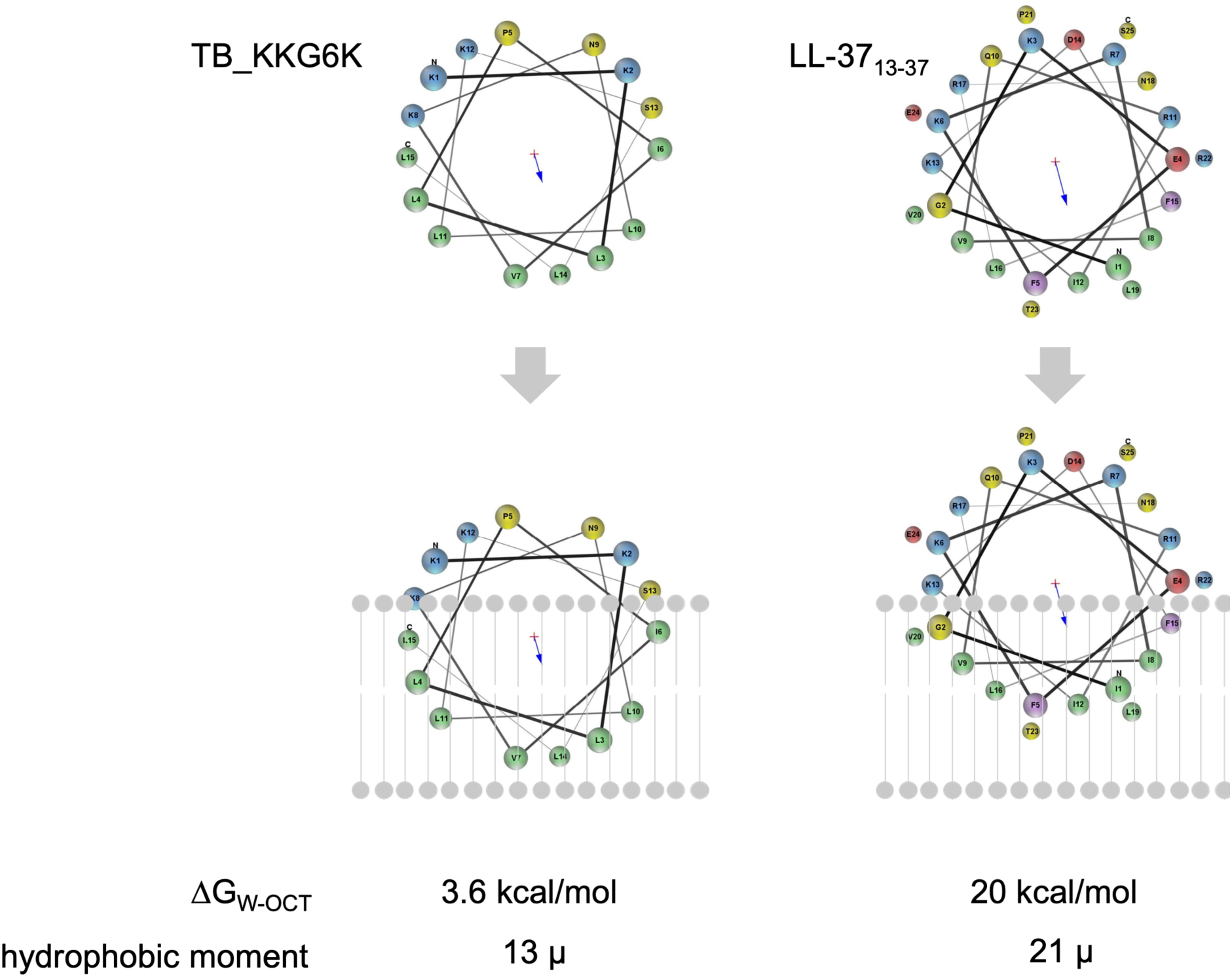
*In silico* evaluation of peptide-membrane binding. (The helical wheel projections of TB_KKG6K and LL-37_13-37_ (reference peptide) were performed with Membrane Protein Explorer (mPEX; 31). The proposed partitioning of each peptide into the phospholipid bilayer (scheme in grey) is shown. The bilayer partitioning free energy (ΔG_W-OCT_) is indicated in kcal/mol and the hydrophobic moment (extent of amphipathicity) is indicated by µ for both peptides. Negatively charged residues are shown in red, positively charged in blue, aliphatic in green, polar in yellow, and aromatic in violet.

### TB_KKG6K rapidly enters *C. albicans* cells

After having identified TB_KKG6K to be a membrane-active peptide in *C. albicans*, we wanted to track its cellular localization and to assess whether it is retained in the cell membrane or enters the target cell. To this end, we used FITC-TB_KKG6K, which had been confirmed to have the same anti-*Candida* activity as the unlabeled peptide (data not shown). Before treatment with the FITC-labeled TB analog, *C. albicans* cells were stained with 0.8 µM of the fluorescent styryl dye *N*-(3-triethylammoniumpropyl)-4-(6-(4-(diethylamino) phenyl) hexatrienyl) pyridinium dibromide (FM4-64) to observe its membrane distribution by laser scanning microscopy (LSM) (Figure 5). FM4-64 first interacted with the cell membrane, which appeared brightly stained. Over time, the signal faded from the cell membrane due to the dye’s endocytic transport into the cells where it stained intracellular membranes (Figure 5A). We then assessed the localization of 0.5 µM, 1 µM and 2 µM FITC-TB_KKG6K in the presence of FM4-64. Immediately after addition of the peptide at the sub-inhibitory concentrations of 0.5 µM or 1 µM to the cells, the peptide-specific FITC signal localized at the cell membrane and accumulated intracellularly in round compartments, presumably vacuoles. In some cells, however, the fluorescence appeared dispersed throughout the cytoplasm (Figure 5B). The intensity of the cytoplasmic fluorescent signal dramatically increased when the cells were exposed to 2 µM FITC-TB_KKG6K, which corresponded to its IC_90_ (Figure 5B). Furthermore, the number of peptide-containing cells increased in proportion to the applied peptide concentration, and at 2 µM, the peptide had entered almost all of the cells in the focal plane (Supplementary Figure S1). Of note, in the presence of 0.5 µM or 1 µM peptide, FM4-64 fluorescence was not retained in the cellular membranes as was the case in the control cells (Figure 5A), rather, the dye accumulated in presumptive pre-vacuolar compartments or was found dispersed throughout the cytoplasm (Figure 5B). With 2 µM FITC-TB_KKG6K treatment, the FM4-64 fluorescence signal had dissipated throughout the whole cell where it co-localized with that of the peptide (Figure 5B, Supplementary Figure S1). This observation suggested that the peptide’s activity disturbed the membrane accumulation of FM4-64 and further underlines that TB_KKG6K affects the integrity of the cell membrane and the intracellular membranes. No retention of the peptide in the cell wall was observed (Supplementary Figure S2A, B).

**Figure 5.**
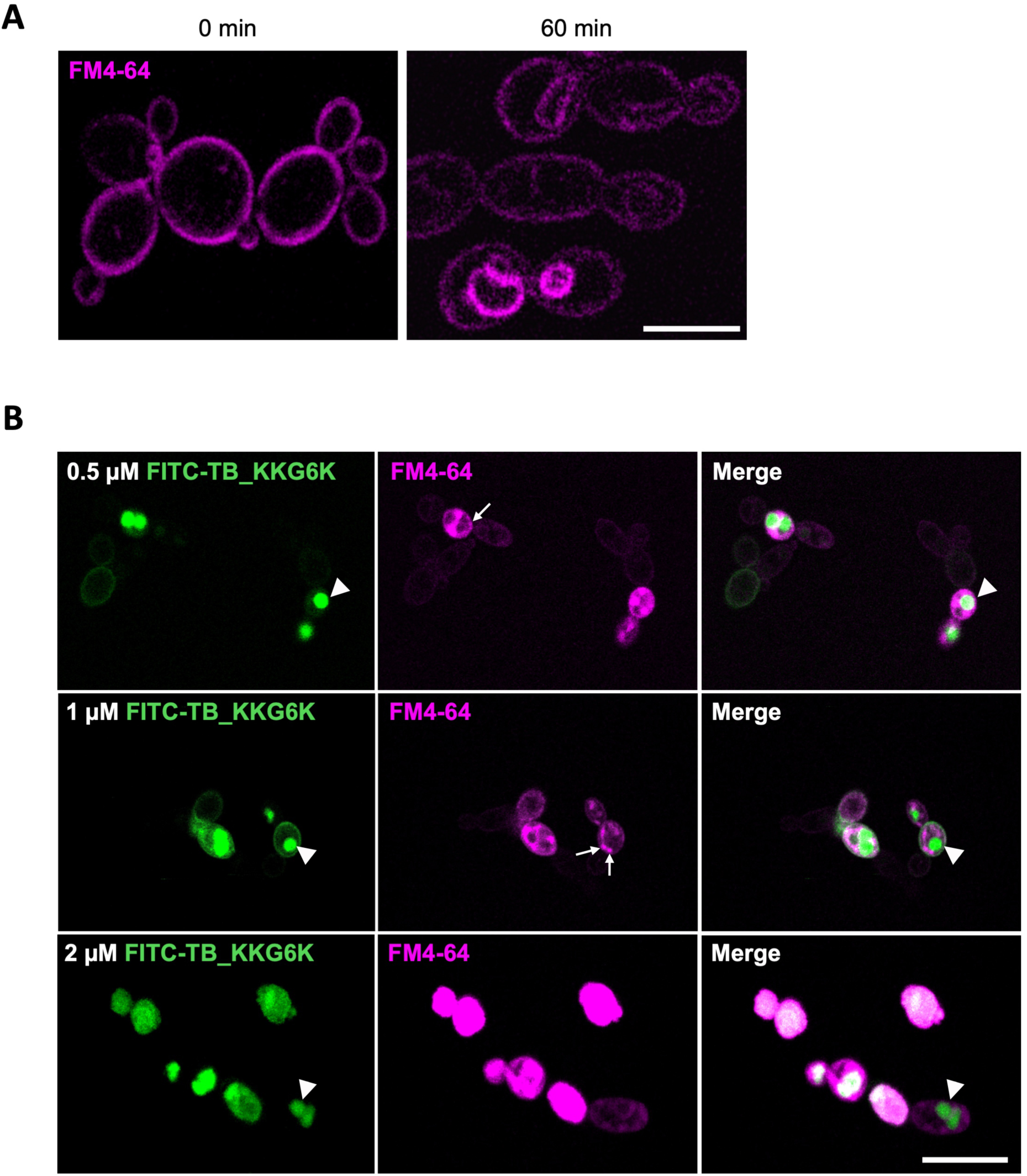
Laser scanning microscopic imaging of *C. albicans* stained with FM4-64 and treated with FITC-labelled TB_KKG6K in *C. albicans*. (A) The binding of 0.8 µM FM4-64 to the cell membrane and intracellular membranes was monitored in *C. albicans* incubated for 0 min and 60 min at 30°C. (B) Cells were exposed to 0.5 µM, 1 µM and 2 µM peptide in the presence of 0.8 µM FM4-64 for 5 min at 30°C. Presumptive vacuoles and pre-vacuolar compartments are marked with arrowheads and small arrows, respectively. Scale bar, (A) 5 µm; (B) 10 µm.

The extremely fast interaction of the peptide with the *C. albicans* cells was further confirmed by fluorescence-activated cell sorting (FACS) analysis. The relative amount of FITC-positive cells was quantified after 5 min and 30 min of peptide exposure. After only 5 min of incubation, 89.7±8.6% of the 5,000 counted cells in total were identified to be FITC-positive. The number of cells that had interacted with the peptide did increase only slightly with time, reaching 91±6.9% after 30 min of incubation. This indicates that the peptide rapidly interacts with *Candida* cells, confirming the measurements on membrane activity and our microscopic observations.

The distribution of the TB analog in the cytoplasm raised the question how the peptide enters the yeast cell. We, therefore, exposed *C. albicans* to FITC-TB_KKG6K at its inhibitory concentration, applying conditions that reduce respiration and cellular metabolism, the aim being to discriminate between passive translocation and active, endocytic cell entry of the peptide. The yeast cells were incubated for 15 min with 2 µM of the peptide, either at 4°C, or at 30°C in the presence of 100 µM carbonyl cyanide m-chlorophenylhydrazone (CCCP), a chemical inhibitor of oxidative phosphorylation. Irrespective of the experimental setting, the peptide was always localized in the cytoplasm, similarly to cells incubated with the peptide at standard conditions (at 30°C without CCCP) (Figure 6).

**Figure 6.**
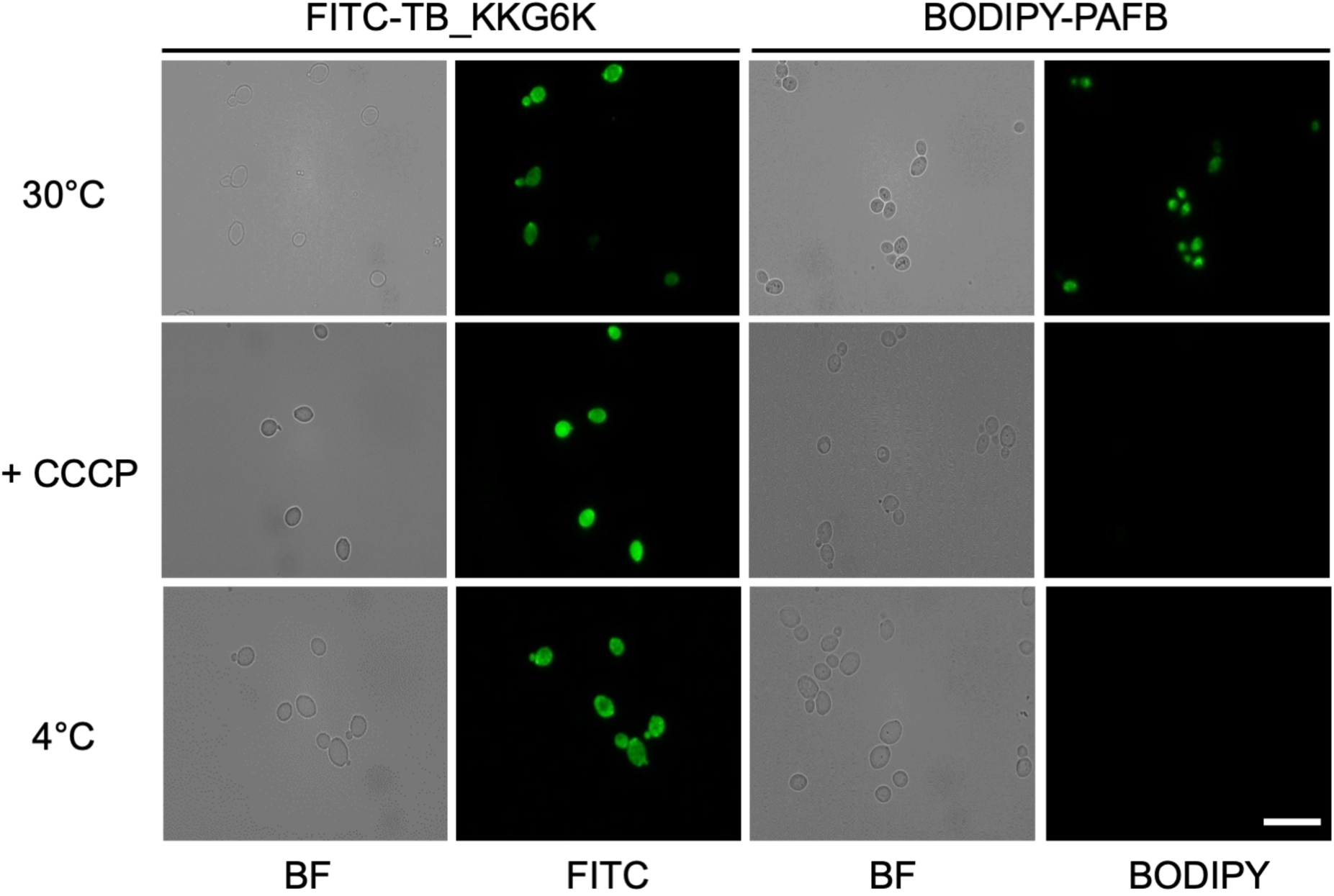
TB_KKG6K entry into *C. albicans* cells. Cells were exposed to 2 µM of FITC-TB_KKG6K for 15 min at 30°C (standard condition), at 30°C in the presence of 100 µM CCCP (inhibition of respiration) or at 4°C (reduction of cellular metabolism) before fixation. As a control for energy-dependent uptake, *C. albicans* was incubated with 2 µM BODIPY-PAFB for 45 min under the same incubation conditions as described above and then fixed for fluorescence microscopy. BF, brightfield; FITC, fluorescence microscopy. Scale bar, 15 µm.

For comparison, we applied the *P. chrysogenum* antifungal protein B (PAFB), which was previously shown to require energy and an active cellular metabolism to enter *C. albicans* cells (32). PAFB was conjugated with the green fluorophore 4,4-difluoro-5,7-dimethyl-4-bora-3a,4a-diaza-s-indacene-3-propionyl ethylenediamine hydrochloride (BODIPY) and incubated with the yeast cells at 30°C for 45 min. As expected, the labeled PAFB (BODIPY-PAFB) was taken up by *C. albicans* under these experimental conditions, but no internalization was observed in cells that had been exposed to the BODIPY-PAFB at 4°C or in the presence of CCCP at 30°C (Figure 6), proving that PAFB requires energy and active metabolism for cell entry.

This suggests that TB_KKG6K, unlike PAFB, enters the yeast cells in an energy-independent manner when applied at its effective concentration.

### TB_KKG6K disintegrates subcellular structures in *C. albicans*

Knowing that TB_KKG6K acts in a fungicidal way (7) and having localized the peptide in the cytoplasm of exposed cells, we performed electron microscopy of cryo-fixed cells to get an idea about the cellular damage induced in *C. albicans* by the application of a high concentration of the peptide. The submicroscopic architecture of the majority of untreated cells appeared intact (Figure 7A, B), with all components known from budding yeasts (33, Figure 3). Nuclei, mitochondria, strongly stained vacuoles, late endosomes (multi-vesicular bodies: MVB), endoplasmic reticulum (ER), ribosomes and glycogen were constantly observed; peroxisomes (microbodies), elements of the inconspicuous Golgi apparatus, and small, endocytic/exocytic vesicles were present as well, albeit at a lower frequency. The exposure to TB_KKG6K caused severe ultrastructural damage to the vast majority of cells (Figure 7C-G), except for a few morphologically intact individuals (Figure 7C, arrows). The most remarkable features, for example, were (i) heavy mitochondrial damage to the extent that those organelles (and their characteristic membrane architecture) were eventually hard to identify (Figure 7D-G), (ii) disrupted nuclear envelope (Figure 7F), (iii) increased frequency of late endosomes (MVB: Figure 7D), and (iv) strongly stained, partly irregularly shaped patches of approximately 45-60 nm in width. In some cases, it was possible to identify such patches as membrane-bound, single or clustered vesicles, indicating endocytic traffic (Figure 7D); some patches likely represented glycogen rosettes, some possibly aggregated polyribosomes, however, further work is needed for a more detailed characterization. Notably, the cellular shape of peptide-treated *C. albicans* along with the structure of its cell membrane and cell wall remained generally intact, and indications of cell lysis were not observed (Figure 7C-G), with the possible exception of very sporadic, local cell membrane rupture (data not shown).

**Figure 7.**
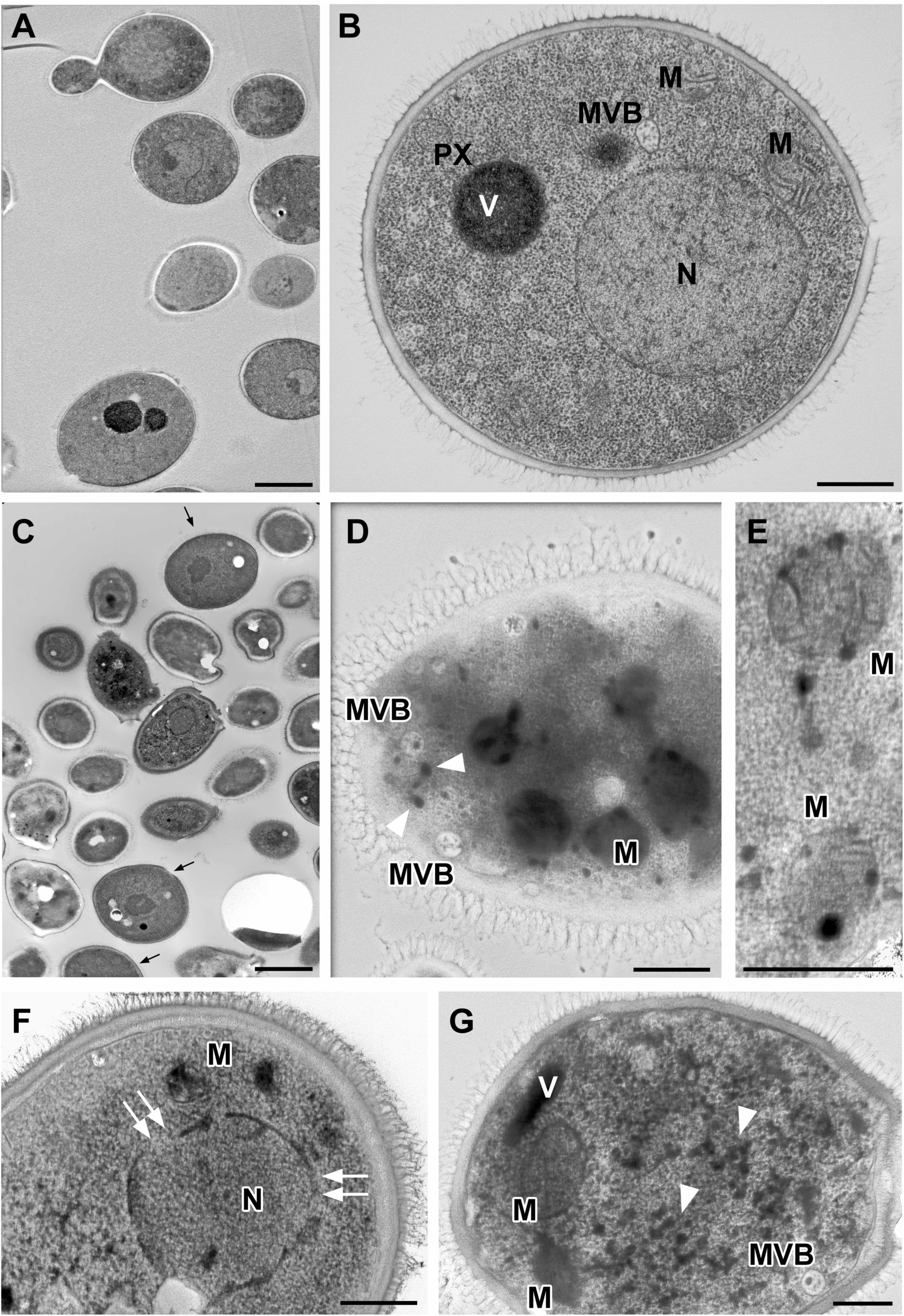
Changes in the cellular morphology of *C. albicans* in response to TB_KKG6K treatment. Electron micrographs of cryo-fixed *C. albicans* cells left untreated (controls, panels A, B) or exposed to 30 µM TB_KKG6K (panels C-G) at 30°C for 60 min. Scale bars, 2 µm (in panels A, C) and 500 nm (in panels B, D, E, F, G), respectively. (A,B) Normal ultrastructure in untreated *C. albicans* serving as controls with nucleus (N), mitochondria (M), late endosomes (multi-vesicular bodies: MVB), vacuoles (V), peroxisomes (P) and abundant ribosomes and glycogen (not specifically highlighted) throughout the cytoplasm (ER and Golgi not depicted in those section planes). (C-G) Various patterns of subcellular degradation in TB analog treated cells. (C) Overview showing only a few, visually intact cells (arrows). (D) Disintegrated mitochondria (M) are hardly recognizable in contrast to numerous MVBs and strongly stained, small vesicles about 55 nm in width (arrowheads). (E) Damaged, though recognizable mitochondria (M) with still visible membrane remnants. (F) Ruptured envelope (double arrows) of the damaged nucleus (N). (G) Disintegrated mitochondria (M), MVB and a vacuole (V), as well as strongly stained, about 55 nm-wide patches (arrowheads), presumably (endocytic) vesicles and/or glycogen rosettes and/or ribosomal aggregates.

## Discussion

Recently, we reported for the first time that TB_KKG6K acts in a fungicidal manner against the opportunistic human pathogenic yeast *C. albicans* (7). Previous studies on TB peptides and TB derivatives confirmed the ability of those compounds to inhibit bacterial growth due to their membrane perturbing activity (9, 10, 34). The membrane activity of AMPs is hypothesized to play a significant role in the induction of expeditious killing of microbial cells (12, 35–37). Based on the high antifungal efficacy of TB_KKG6K at killing planktonic and sessile *C. albicans* cells *in vitro* (7), we focused, in the present study, on obtaining insights into the antifungal mechanism of action of this TB analog.

Using fluorescent dyes for the assessment of membrane depolarization (DiSC_3_(5)) and permeabilization (SYTOX Green), we could show that TB_KKG6K impaired cell membrane integrity. The TB analog induced an immediate depolarization and permeabilization of the fungal cell membrane in a concentration and time dependent manner (Figure 1, Figure 2). The rapid reaction at the cell membrane does not allow any conclusion as to which of these two events precedes the other. Neither membrane depolarization nor permeabilisation is *per se* lethal as long as the cell is able to restore cell membrane function, e.g. under exposure to sub-lethal peptide concentrations (38). Cell membrane depolarization, however, often coincides with cell membrane permeabilization, resulting in severe membrane perturbation with excessive loss of ions and metabolite gradients due to membrane leakage or pore formation, affecting cell viability or even killing the cell by membrane rupture (39, 40). Persistent pores in the cell membrane and leakage of the cell content in response to exposure to TB_KKG6K, however, were not observed (Figure 5, Figure 7). Therefore, the TB analog-induced cell membrane permeability is likely to be altered by the establishment of transient pores that are large enough to allow the nucleic acid binding dye SYTOX Green to gain access into the cell (41–43), but too small to be detectable by electron microscopy (Figure 7). In this regard, also the leakage of the low molecular weight compound ANTS/DPX from LUVs that had been exposed to the TB analog is presumed to be the result of permeabilization but not rupture of the model membranes (44).

The use of model membranes containing different phospholipids and the quantification of the amount of ANTS/DPX dye leaking out from LUVs upon exposure to the TB analog revealed that TB_KKG6K induced leakage in LUVs composed of anionic PG (Table 1). This phospholipid represents the main component of membranes of Gram-positive bacteria (22). This was in agreement with the documented toxicity of the TB analog towards bacterial cells (10). However, the peptide also permeabilized LUVs containing the neutral lipid PC, proving that hydrophobic interactions play a role in its membrane activity. Neutral PC can be regarded as a primary building block of eukaryotic membranes, being one of the most abundant phospholipids in their lipid bilayer (26, 27). When comparing the effect of the TB analog on PC-LUVs with vesicles containing PC in combination with other specific membrane lipids, the leakage was significantly increased in the presence of PI, indicating that this anionic phospholipid may be particularly amenable to TB_KKG6K attack (Table 1). Indeed, fungal cell membranes are generally more negatively charged than human ones due to a higher content of anionic phospholipids such as PI and PA (12). This correlates with the results obtained from the peptide-lipid overlay experiments (Figure 3). TB_KKG6K showed a preference for unphosphorylated PI and PI mono-, bi- and tri-phosphates (Figure 3), which are known to play a crucial role in a number of cell membrane functions, including cell signaling and regulation of protein activity in and at the cell membrane (45, 46). Of particular significance is the negative charge of these phospholipids, which specifically attracts cationic AMPs (13, 47). Another membrane lipid that was recognized by the TB analog was CL, an integral mitochondrial membrane component (48, 49). Its combination with PC resulted in increased LUV leakage at the highest peptide concentration tested (Table 1). This implies that the peptide’s activity is not restricted to the cell membrane, but extends to intracellular targets; in this case, the mitochondrial envelope (50). Interestingly, in comparison to its effects on the membrane composed of only PC, TB_KKG6K evoked significantly lower leakage in PC-LUVs containing ergosterol or sphingomyelin, which are two lipids enriched in fungal (12, 51) and mammalian lipid rafts (52–54), respectively. It has been reported that sterols, including ergosterol, reduced the membrane binding of AMPs such as LL-37 and Temporin L due to the sterol’s ability to cause increased acyl chain order and lipid packing, resulting in an increased membrane density and thickness (55). This might render membrane penetration more difficult for AMPs, thereby conferring a certain degree of membrane protection depending on the type of sterol present. The effect of sphingomyelin on AMP-induced membrane perturbation is comparatively less characterized; however, in line with our observation, the presence of sphingomyelin in model membranes reduced the membrane intercalation of LL-37, both alone and in combination with ergosterol (56). This, coupled with the fact that mammalian cell membranes are generally less negatively charged than fungal membranes, could also explain why TB_KKG6K is well tolerated by primary human cells *in vitro* (7).

Physicochemical investigations on a number of AMPs and cell penetrating peptides have demonstrated that apart from charge and amphipathicity, their secondary structure and conformation play a crucial role in their ability to interact with cellular membranes (57–59). In particular, circular dichroism spectroscopy studies revealed a characteristic shift from a random, disordered secondary structure of TB_KKG6K in an aqueous environment to an α-helical conformation in the presence of sodium dodecyl sulfate, a fungal cell membrane surface mimetic (7). This structural transition is hypothesized to facilitate binding and insertion of peptides into the fungal cell membrane (57, 60, 61). The *in silico* helical-wheel projection studies underlined these data (Figure 4). Compared to the membrane-active anti-*Candida* peptide LL-37_13-37_ (28), the predicted α-helix of TB_KKG6K shows greater potential for its insertion into the hydrophobic portion of the membrane bilayer (Figure 4). This model is further supported by the calculated membrane partitioning free energy and the hydrophobic moment, which are both lower for the TB analog compared to that of LL-37_13-37_ (Figure 4).

The conclusion that TB_KKG6K perturbs the fungal cell membrane and, in terms of energetics, can penetrate the cell membrane with relative ease, was proven by fluorescence microscopy applying a FITC-labelled TB_KKG6K. Co-staining with the lipophilic dye FM4-64 (62) helped follow the entry of the peptide into the fungal cell (Figure 5). The exposure to FITC-TB_KKG6K at its IC_90_ value, however, immediately impaired cell membrane integrity and disrupted the specific membrane binding capability of FM4-64, causing the lipophilic dye to spread across the cell and co-localize with the peptide-specific fluorescent signal in the cytoplasm (Figure 5). Overview images confirmed that only those cells that had internalized the peptide presented a diffuse FM4-64 staining pattern (Supplementary Figure S1). The peptide action, however, was too rapid to capture the presumed initial co-localization of FITC-TB_KKG6K with FM4-64 in the cell membrane prior to peptide associated cell membrane perturbation. The swift manner with which the peptide interacted with over 89% of the cells within a very short time of exposure was confirmed by FACS-based quantification of FITC-positive cells. In contrast, the membrane affinity and distinct compartmentalization in presumptive vacuoles of FITC-TB_KKG6K were occasionally visible in the cells that had been exposed to sub-inhibitory peptide concentrations (Figure 5). Several possible explanations might account for this observation. (i) Peptide entry into the cell is concentration dependent, as has been shown for other AMPs like the hexapeptide PAF26 (63), Histatin-5 (64), or pVEC (65). At sub-inhibitory concentrations, these peptides are internalized by receptor-mediated endocytosis and accumulate in the vacuole, while they directly translocate into the cytoplasm of the target cell at their effective concentration, inducing cell death. (ii) The concentration of the TB analog has no influence on the mode of internalization, and cellular entry always occurs in an energy independent manner. Interestingly, Maniti et al. describe a hypothesized ‘energy-independent endocytic process’ that allows the cell penetrating peptide penetratin to induce membrane curvature in artificial lipid membranes due to its positive charge. If the angle of curvature crosses a certain threshold, it could cause the membrane to fold back upon itself, forming a peptide containing, endosome like invagination and facilitating peptide uptake independent of energy (66). If the mode of TB_KKG6K entry into the yeast cell took place in a similar manner, the uptake of the peptide at sub-inhibitory concentrations and its dispersion from the vacuole into the cytoplasm would occur slower than at the effective concentration, which would allow the vacuolar localization of the peptide to be observed microscopically. The question how TB_KKG6K enters the fungal cell certainly merits further investigations in the future. In contrast to PAFB, however, which is internalized *via* endocytosis (32), the entry of TB_KKG6K into *C. albicans* cells at the peptide’s effective concentration was hampered neither by the reduction of cellular metabolism nor the inhibition of oxidative phosphorylation, excluding the involvement of a receptor-mediated endocytic pathway at this concentration (Figure 6). Passive entry into microbial cells is considered more advantageous in terms of antimicrobial efficacy, as energy dependent AMPs easily lose their activity under physiological conditions of reduced metabolic activity, e.g. in cells growing in a biofilm (67–69). The accumulation of FITC-TB_KKG6K in the vacuole, when applied at sub-inhibitory peptide concentrations, might also protect the cells from peptide induced cell death. In fact, a similar phenomenon was observed with many AMPs (12, 63, 64), including the *P. chrysogenum* antifungal proteins PAF, PAFB and PAFC (70, 71). The induction of cell death was closely linked with the cytoplasmic localization of the AMPs, but did not take place as long as the peptides resided in the vacuole. Either these peptides are degraded, or they overcome vacuolar function; with the latter scenario, it is then only a matter of time until they disrupt the vacuolar membrane, spread into the cytoplasm and kill the cell.

Finally, transmission electron microscopy revealed a severe disintegration of intracellular structures in response to the treatment of *C. albicans* cells with TB_KKG6K (Figure 7). Although the cell content was heavily damaged, the overall cell shape was retained and the structure of the cell wall and the cell membrane showed no signs of rupture in the presence of the TB analog (Figure 7). Since TB_KKG6K translocates into the cell and intracellular membranes contain high amounts of anionic phospholipids and PIPs, it is reasonable to assume that the peptide interacts also with the subcellular membranes, damaging the respective organelles, e.g. mitochondria, nuclei (Figure 7). The ability of TB_KKG6K to permeabilize model membranes containing CL (Table 1) further suggests that the peptide interacts with the CL that resides in the inner mitochondrial membrane. CL is required for the maintenance of respiration and the regulation of reactive oxidative species (ROS) generation (48). Impairment of CL function and/or mitochondrial damage by the TB analog could be one reason for the observed induction of intracellular ROS (7), which is known to damage cellular organelles and molecules, including nucleic acids, proteins, and lipids (13, 72).

## Conclusion

Our study provides a rationale behind the rapid killing of *C. albicans* by TB_KKG6K. Based on our results, we propose the following model to explain its mechanism of action: The TB analog preferentially affects membranes composed of anionic but also neutral phospholipids. Due to its physicochemical features (charge, hydrophobicity), we assume that TB_KKG6K binds to the yeast cell membrane *via* hydrophobic interactions, whereby additional electrostatic interactions with anionic membrane lipids stabilizes the peptide lipid binding. Upon cell membrane binding, the peptide impairs the integrity of the cell membrane. Membrane depolarization and permeabilization might facilitate the peptide’s translocation through the membrane into the cell where it interacts with intracellular targets and executes its killing activity (73). The translocation of the TB analog into the yeast cell, its intracellular distribution and the disintegration of subcellular structures but not that of the cell wall or the cell membrane further underlines that TB_KKG6K is a membrane-active, non-lytic peptide that also affects the membrane integrity of intracellular compartments, possibly in combination with the induction of intracellular ROS. The documented membrane activity renders this peptide a promising candidate for the development of next generation anti-*Candida* therapeutics. Due to its rapid, fungicidal activity, this amphibian peptide analog may hamper the onset of drug resistance, which has become increasingly challenging in the treatment of mycoses caused by the opportunistic human pathogen *C. albicans*.

## Materials and Methods

### Microorganisms, media and growth conditions

Single, fresh colonies of *C. albicans* (CBS 5982) grown on solid yeast extract peptone dextrose agar (yeast extract peptone dextrose broth (YPDB with 2% (w/v) agar; Sigma-Aldrich, St. Louis, MO, USA) were used to inoculate 10 mL of YPDB (Sigma-Aldrich, St. Louis, MO, USA). After overnight cultivation at 37°C and shaking at 200 rpm, the cells were counted and diluted in 5% (w/v) PDB (0.05×PDB) to the cell number applied in the respective experiments described below.

### Peptide synthesis and protein expression

TB_KKG6K, and the PCγ^C-terminal^ (amino acid sequence: CGGASCRG; MW 709.8 Da; 14) were synthesized on solid phase, using standard protocols for the Fmoc chemistry and were then purified by reversed-phase high performance liquid chromatography (RP-HPLC) and analyzed by electrospray ionization-mass spectrometry (ESI-MS) as described previously (7, 14). For fluorescence based analyses, the TB analog was synthesized and conjugated to the 6-aminohexanoic acid linker first and then to the green fluorophore fluorescein isothiocyanate (FITC) as described (74). The FITC-conjugated peptide (FITC-TB_KKG6K) was purified by RP-HPLC on a Jupiter 10μ Proteo 90A° (100×21, 20 mm) column, flow rate 20.0 mL/min and analyzed by ESI-MS as described by Kakar et al. (7). Calculated mass (Da): 2221.28; found: 1111.88[M+2H]^2+^; 741.82 [M+3H]^3+^. The *Penicillium chrysogenum* antifungal protein PAFB was expressed using a *P. chrysogenum*-based expression system (75) and purified by cation-exchange chromatography as previously described (32). For fluorescence based analyses, PAFB was labelled with the green fluorophore 4,4-difluoro-5,7-dimethyl-4-bora-3a,4a-diaza-s-indacene-3-propionyl ethylenediamine hydrochloride (BODIPY™ FL EDA, Invitrogen, Waltham, MA, USA) as described (32).

### Broth microdilution assays

The IC_90_ of the TB analog, octenidine (N,Nʹ-(1,10 decanediyldi-1[4H]-pyridinyl-4-ylidene)-bis-(1-octanamine) dihydrochloride), Schülke and Mayr GmbH, Vienna, Austria), and PCγ^C-terminal^ (14) was determined for *C. albicans* using broth microdilution assays carried out in 96-well microtiter plates (Nunclon Delta, Thermo Fisher Scientific, Waltham, MA, USA). The assay was performed with 1×10^4^ cells/mL and 1×10^6^ cells/mL in 0.05×PDB. Briefly, 100 µL of either cell concentration were mixed with 100 µL of two-fold compound dilutions prepared in 0.05×PDB in a 96-well microtiter plate. The final concentration range of the compounds tested was 0-32 µM. The plates were incubated at 30 °C for 24 h under static conditions. The cells were then resuspended by vigorous pipetting, and the optical density (OD) of the resuspended cell suspension was measured with a multimode microplate reader (FLUOstar Omega, BMG Labtech, Ortenberg, Germany) at a wavelength of 620 nm. The OD_620_ of the untreated control was assigned 100% growth. All samples were prepared in technical triplicates and the assays were repeated at least twice.

### Membrane depolarization assay

*C. albicans* cells were prepared in 0.05×PDB at a concentration of 1×10^6^ cells/mL and 100 μL was added to each well of a black polystyrene microtiter plate (Greiner Bio-One, Frickenhausen, Germany). Fluorescence intensity was followed for 7 min 30 sec at an excitation/emission wavelength of 622/670 nm using a multimode microplate reader (CLARIOstar Plus, BMG Labtech, Ortenberg, Germany) to obtain baseline values for cell and medium background fluorescence. Measurements of the fluorescence intensity were then paused, and 50 μL of the 3,3′-dipropylthiadicarbocyanine iodide (DiSC_3_(5)) dye (Thermo Fisher Scientific, Waltham, MA, USA), prepared separately in 0.05×PDB at a concentration of 6 µM, was added to each well. Fluorescence measurements were continued until a stable signal was achieved. Serial dilutions of the TB analog, Triton X-100 (positive control) and PCγ^C-terminal^ (negative control) were prepared in 0.05×PDB in a separate 96-well microtiter plate (Nunclon Delta, Thermo Fisher Scientific, Waltham, MA, USA) in a final volume of 50 μL per well. For the untreated control, 50 μL of 0.05×PDB replaced compound addition. Measurements of the fluorescence intensity were paused, and 50 µL of the serially diluted compounds were added to the cell suspension, resulting in a final volume of 200 μL per well and a final DiSC_3_(5) concentration of 1.5 μM. The final concentrations of the positive and negative control were 1% (w/v) and 32 μM, respectively. The TB analog was tested at a final concentration range of 0-2 μM.The fluorescence intensity measurements were recommenced after compound addition and were recorded at 30°C in 5 min intervals for a further 60 min. The fluorescence values obtained post compound addition were background corrected (samples without cells) and normalized by subtracting the values of the untreated control from the values of the treated samples.

### SYTOX Green uptake assay

*C. albicans* cells were prepared in 0.05×PDB at a concentration of 1×10^6^ cells/mL. The SYTOX Green nucleic acid stain (Thermo Fisher Scientific, Waltham, MA, USA) was then added to the cell suspension at a final concentration of 0.2 µM. The cells were incubated in the dark at 30°C for 5 min under static conditions. In the meantime, serial dilutions of the TB analog, the positive control octenidine, and the negative control PCγ^C-terminal^ were prepared in 0.05×PDB in a 96-well microtiter plate (Nunclon Delta, Thermo Fisher Scientific, Waltham, MA, USA) at a volume of 100 μL per well. The cell suspension, after pre-incubation with SYTOX Green, was distributed in 100 μL aliquots per well in the compound containing 96-well microtiter plate (Nunclon Delta, Thermo Fisher Scientific, Waltham, MA, USA). The final concentrations of the positive and negative controls were 2 μM and 32 μM, respectively. The TB analog was tested at a final concentration range of 0-2 μM. For the untreated control, 0.05×PDB replaced compound addition. Fluorescence intensities were measured in 5 min intervals for 60 min at 30°C using an excitation/emission wavelength of 480/530 nm in a multimode microplate reader (CLARIOstar Plus, BMG Labtech, Ortenberg, Germany). The fluorescence values were background corrected and normalized by subtracting the values of the untreated control from the values of the treated samples.

### Phosphoinositide binding assay

The lipid-peptide overlay assay was performed using FITC-conjugated TB_KKG6K and PIP strips according to the manufacturer’s instructions (Echelon Biosciences, Salt Lake City, UT, USA). To ensure the suitability of the applied detection method, 0.5 µg of FITC-TB_KKG6K was directly dotted in a volume of 1 µL onto the PIP strip membranes. The dotted peptide was allowed to dry for 30 min at 30°C before the PIP strips were tested with FITC-TB_KKG6K. Briefly, the PIP strips were blocked in blocking buffer (10 mM Tris [pH 8.0]; 150 mM NaCl; 0.1% Tween 20; 3% fatty acid-free bovine serum albumin (BSA), Sigma-Aldrich, St. Louis, MO, USA) for 1 h. All of the subsequent experimental procedures were performed in the dark.

The PIP strips were incubated for 1 h with FITC-TB_KKG6K and diluted in blocking buffer to a final concentration of 1.5 µg/mL. Then, the membranes were washed 3 × 10 min in blocking buffer. The FITC-TB_KKG6K bound to specific phosphoinositides was detected fluorometrically with a Typhoon FLA 9500 biomolecular imager (GE Healthcare, Chicago, IL, USA) equipped with a 473 nm laser and filters for excitation/emission wavelengths of 494/520 nm. The FITC fluorescence signal intensity was semi-quantified by the ImageJ/FIJI software (version 1.8.0/1.53q, U.S. National Institutes of Health, Bethesda, MD, USA).

### Vesicle leakage assay

For the leakage experiments, the following lipids were used: 1-palmitoyl-2-oleoyl-glycero-3-phosphocholine (POPC), 1-palmitoyl-2-oleoyl-sn-glycero-3-phosphoethanolamine (POPE), 1’,3’-bis[1,2-dioleoyl-sn-glycero-3-phospho]-glycerol, sodium salt (cardiolipin, CL), egg sphingomyelin, 1-palmitoyl-2-oleoyl-sn-glycero-3-(phospho-rac-(1-glycerol)) (POPG), 1-palmitoyl-2-oleoyl-sn-glycero-3-phosphoinositol, ammonium salt (POPI), 1-palmitoyl-2-oleoyl-sn-glycero-3-phospho-L-serine, sodium salt (POPS), ergosterol, N-palmitoyl-D-erythro-sphingosine (ceramide). All lipids were purchased from Avanti Polar Lipids (Alabaster, AL, USA).

Lipid films composed of POPC, POPG, POPC/POPI (9:1 mol), POPC/POPS (9:1 mol), POPC/POPE (9:1 mol), POPE/POPG/CL (6:2:1 mol), POPC/ergosterol (4:1 mol), POPC/Ceramide (9:1 mol), POPC/CL (3:1 mol), POPC/sphingomyelin (9:1 mol), POPC/POPE/ergosterol ([POPC:POPE 9:1 mol]; [POPC:POPE]:ergosterol 4:1 mol), POPC/POPE/ergosterol/ceramide (56:6:8:20 mol), and POPC/POPS/ergosterol ([POPC:POPS 9:1 mol];[POPC:POPS]:ergosterol 4:1 mol) were prepared by dissolving the appropriate lipids, prepared in glass tubes at a total amount of 20 mg, in chloroform/methanol (2:1 v/v). The glass tubes were subsequently placed under a stream of nitrogen to allow the solvent to evaporate. They were then stored in vacuum overnight to completely remove all residual traces of solvent. The prepared lipid films were hydrated in 10 mM HEPES buffer (pH 7.4) containing 68 mM NaCl, 12.5 mM 8-aminonaphthalene-1,3,6-trisulfonic acid (ANTS; Molecular Probes, Eugene, ORE, USA) and 45 mM p-xylene-bis-pyridinium bromide (DPX; Molecular Probes, Eugene, ORE, USA). The hydration process was carried out with intermittent vortexing for 1 min at the time points 0, 5, 10, 20, 30, and 60 min. For each formulation, a temperature well above the gel-to-fluid phase transition of its constituent lipids was applied. The hydrated lipid films were then extruded 25 times using a handheld mini extruder (Avanti Polar Lipids, Alabaster, AL, USA) with a 100 nm pore diameter polycarbonate filter to obtain 100 nm LUVs. Unilamellarity and size were confirmed by dynamic light scattering (DLS) using a Zetasizer Nano (ZSP, Malvern Panalytical, Prager Electronics, Wolkersdorf, Austria).

The ANTS/DPX containing LUVs were separated from free ANTS/DPX by exclusion chromatography using a column filled with Sephadex™ G-75 (Amersham Biosciences, Amersham, United Kingdom) fine gel swollen in an isosmotic buffer (10 mM HEPES, 140 mM NaCl, pH 7.4). After the void volume, fractions were collected and the phospholipid concentration was determined by phosphate analysis as described by Piller et al. (76). The LUVs containing ANTS/DPX were stored in the dark at room temperature.

The leakage of aqueous contents from the prepared LUVs induced by the TB analog was determined using the GloMax® Discover Microplate Reader (Promega Corporation, Madison, WI, USA). Briefly, serial dilutions of the peptide were prepared in HEPES buffer (10 mM HEPES, 140 mM NaCl, pH 7.4) in a black 96-well microtiter plate (Nunclon Delta, Thermo Fisher Scientific, Waltham, MA, USA) in a final volume of 10 µL per well. Ninety µL of the respective LUVs was added to each well, resulting in a final lipid concentration of 50 µM in a final volume of 100 µL per well. For the untreated lipid control, 10 µL of HEPES buffer was used instead of the peptide. Ten µL of 10 % (v/v) Triton X-100 was used as the positive control. The peptide PCγ^C-terminal^ was used as a negative control at a final concentration of 32 µM. The leakage measurements were started immediately after LUV addition. Fluorescence spectra were obtained at 37°C over a period of 90 min using an excitation/emission wavelength of 360/530 nm. Data are presented in terms of % leakage and were calculated using Equation (1):

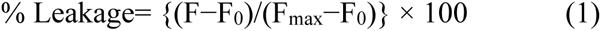

where F is the measured fluorescence, F_0_ is the fluorescence with only the lipid (untreated control) and F_max_ is the fluorescence corresponding to 100 % leakage obtained by the addition of 1 % (v/v) Triton X-100.

### *In silico* evaluation of peptide-membrane binding

Totalizer, a tool provided by Membrane Protein Explorer, mPEX v3.3.0, was used to characterize the binding of TB_KKG6K in comparison to the reference peptide LL-37 (amino acid sequence: LLGDFFRKSKEK**IGKEFKRIVQRIKDFLRNLVPRTES**; MW 4493.3 Da; 28) to the lipid membrane-water interface (31). For the latter, amino acids 13-37, marked in bold, were considered for analysis. *In silico* helical wheel projection, bilayer partitioning free energy (ΔG), and hydrophobic moment (µ) were obtained as described by Piller P et al. (76).

For ΔG, values were calculated using the octanol scale, which calculates the free energy of peptide transfer from water to the hydrophobic region of the membrane bilayer/to octanol.

### Assessment of the cellular localization and uptake mechanism of the TB_KKG6K

To follow the subcellular distribution, two different strategies were pursued using the FITC-conjugated TB_KKG6K. In the first approach, 2 µM FITC-TB_KKG6K was added along with 5 μM of the cell wall specific fluorescent dye calcofluor white (CFW; Sigma-Aldrich, St. Louis, MO, USA) to 800 μL of the *C. albicans* cell suspension (1×10^6^ cells/mL in 0.05×PDB) and incubated with the cells for 15 min at 30°C in the dark to determine if the conjugate localizes with the fungal cell wall. The cell suspension was then washed once with phosphate-buffered saline (PBS; 0.5% (w/v) KH_2_PO_4_, 2.8% (w/v) K_2_HPO_4_, 9% (w/v) NaCl) and fixed in 100 µL of 4% (v/v) formaldehyde (FA; Carl Roth Gmbh & Co., Karlsruhe, Germany) for 10 min. The FA was removed by washing with PBS and 2×10^5^ cells (in 0.05×PDB) per well were transferred to Ibidi^®^ µ-slide 8-well chambered coverslips (Ibidi GmbH, Gräfelfing, Germany) for LSM. The second set of experiments was performed to assess the *in vivo* membrane binding of the peptide and its cellular distribution. To this end, 2×10^5^ *C. albicans* cells (in 0.05×PDB) per well were distributed in Ibidi^®^ µ-slide 8-well chambered coverslips and stained with 0.8 μM of the membrane specific fluorescent styryl dye *N*-(3-triethylammoniumpropyl)-4-(6-(4-(diethylamino) phenyl) hexatrienyl) pyridinium dibromide (FM4-64; Thermo Fisher Scientific, Waltham, MA, USA) in the presence of 0 μM (control), 0.5 μM, 1 μM and 2 μM of FITC-TB_KKG6K. Live cell imaging, performed at 30°C, started immediately after compound addition.

For imaging both approaches, LSM was carried out using an HC PL APO 40×/1.10 CS2 water immersion objective on an SP8 confocal microscope (Leica Microsystems, Wetzlar, Germany), equipped with an 80 MHz pulsed white light laser (WLL) and a 405 nm CW diode laser. HyD detectors were used for optimal fluorescence imaging. Images of CFW (excitation, 405 nm diode; emission, 415 to 479 nm), FITC-TB_KKG6K (excitation, 495 nm WLL; emission, 515 to 530 nm) and FM4-64 (excitation, 515 nm WLL; emission, 670 to 700 nm) were acquired using the Leica Application Suite X (LAS X, version 3.5.7.23225) and further processed by ImageJ/FIJI.

To study the uptake mechanism into the yeast cells, 2 µM FITC-TB_KKG6K was added to 400 μL of a *C. albicans* cell suspension (1×10^6^ cells/mL in 0.05×PDB) and incubated for 15 min at 30°C in the dark. To uncouple mitochondrial oxidative phosphorylation the cells were incubated with 100 µM carbonyl cyanide m-chlorophenylhydrazone (CCCP, Sigma-Aldrich, St. Louis, MO, USA) in the presence of FITC-TB_KKG6K, applying the same incubation conditions. The metabolic activity of the cells was reduced by incubating the cells with FITC-TB_KKG6K at 4°C instead of 30°C. For comparison with an AMP that is taken up by *C. albicans* in an energy-dependent way, PAFB conjugated with the fluorophore BODIPY (BODIPY-PAFB), prepared as described previously (33), was used. In brief, cells were exposed to 2 µM BODIPY-PAFB applying the same experimental conditions as described above. After 45 min of incubation, the cells were washed in PBS and fixed in 100 µL 4% (v/v) FA for 10 min. After washing in PBS, the cells were mounted on glass slides for microscopic analysis. Microscopic imaging of these samples was performed with a fluorescence microscope (Axioplan, Carl Zeiss GmbH, Oberkochen, Germany), equipped with excitation/emission filters (500/535 nm or 450/515 nm for BODIPY or FITC, respectively) and an AxioCam MR3 camera (Carl Zeiss GmbH, Oberkochen, Germany). The images were processed and edited with Axiovision (blue edition), GNU Image Manipulation Program (GIMP, version 2.8.20; www.gimp.org), and Microsoft PowerPoint (Microsoft Corp., Albuquerque, NM, USA) software.

### Ultrastructural analysis of *C. albicans*

For electron microscopy, 3×10^8^ *C. albicans* cells in 1 mL of 0.05×PDB were exposed to 30 µM of TB_KKG6K for 1 h at 30°C under continuous shaking. Untreated cells served as a control. Cell suspensions were slightly pelleted and subjected to rapid cryo-fixation by means of high-pressure freezing and freeze-substitution, followed by epoxy resin embedding essentially as previously described (77), except that the freeze-substitution media contained only 0.8% (w/v) uranyl acetate. Thin sections were optionally poststained with lead salts and viewed with a CM120 transmission electron microscope (Philips, Eindhoven, The Netherlands) equipped with a MORADA digital camera (EMSIS, Münster, Germany).

### Fluorescence-activated cell sorting

*C. albicans* cells were diluted to 4×10^6^ cells/mL in 0.05×PDB and 100 µL of this cell suspension was exposed to 100 µL of FITC-TB_KKG6K prepared in 0.05×PDB to reach a final peptide concentration of 2 µM. The cells were incubated in the dark for 5 min and 30 min, respectively, at 30 °C under static conditions. Untreated cells were used as the negative control. After peptide exposure, the cells were pelleted (900×g for 5 min) and washed twice in PBS. At least 5,000 cells were counted per run and FITC-positive cells were detected with a FlowSight imaging flow cytometer (Amins, Merck Millipore, Billerica, MA, USA) equipped with lasers at 405 nm (violet), 488 nm (blue), and 642 nm (red) wavelengths. The FITC signal was measured utilizing a 488 nm excitation laser and emission in the channel 2 window. Gating was adjusted to reach at least 99% of untreated cells; debris was excluded during data acquisition. For data analysis, the Image Data Exploration and Analysis software (IDEAS; Amins, Millipore, Billerica, MA, USA) was applied. Experiments were repeated three times.

### Statistical analysis

For calculation of significant differences, Two-way ANOVA followed by Dunnett’s multiple comparisons test was performed using GraphPad Prism version 9.1.0 for macOS. *p*-values of ≤0.05 were considered as significant.

## Acknowledgments

F.M., and N.M. designed the experiments and conceived and supervised the study. A.K. performed experiments, A.K., N.M., and F.M. analyzed the data. L.E.S.-V. performed LSM. M.H. performed electron microscopy. L.G. and C.P. performed FACS analysis. J.H. prepared the samples for electron microscopy, A.R. provided TB_KKG6K. G.V. provided PCγ^C-terminal^. A.K. prepared the first manuscript draft. All authors contributed to manuscript writing and revision and approved the submitted version. We thank W. Salvenmoser for helpful discussion, and D. Bratschun-Khan, K. Gutleben and B. Witting for technical assistance. The study was funded by the Austrian Science Fund FWF (HOROS Doctoral Program, W1253 DK HOROS) to F.M. L.G. was financed by the Hungarian National research, Development and Innovation Office - NKFIH, FK 134343 project.

We have no conflict of interest to declare.

