## Supplemental Data for "The membrane activity of the amphibian Temporin B peptide analog TB_KKG6K sheds light on the mechanism that kills *Candida albicans*"

Running Head: Membrane activity of the TB_KKG6K against *C. albicans*

**Keywords:** Temporin B, TB analog, antifungal peptide, *Candida albicans*, membrane activity, depolarization, permeabilization, leakage, uptake

**Figure S1**

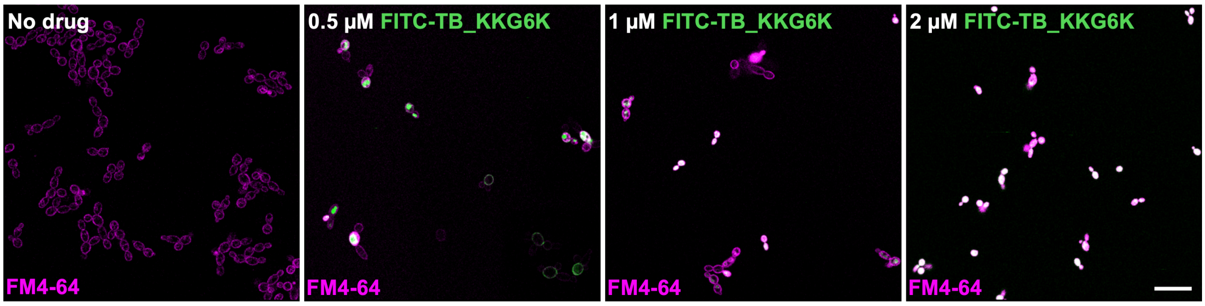


**Figure S1.** Laser scanning microscopic images of *C. albicans* treated with TB_KKG6K. Cells were exposed to 0.8 µM of the lipophilic membrane specific dye FM4-64 in the absence (no drug, control) or presence of 0.5 µM, 1 µM and 2 µM FITC-labelled TB_KKG6K for 15 min at 30°C. Scale bar, 20 µm.

**Figure S2**

**A**

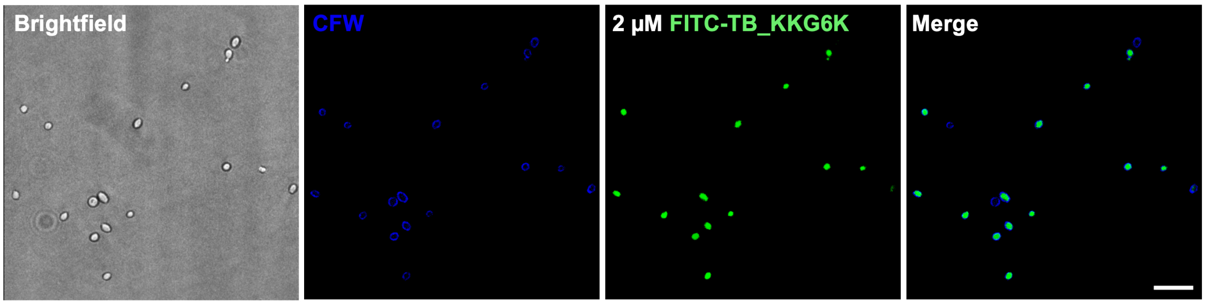


**B**

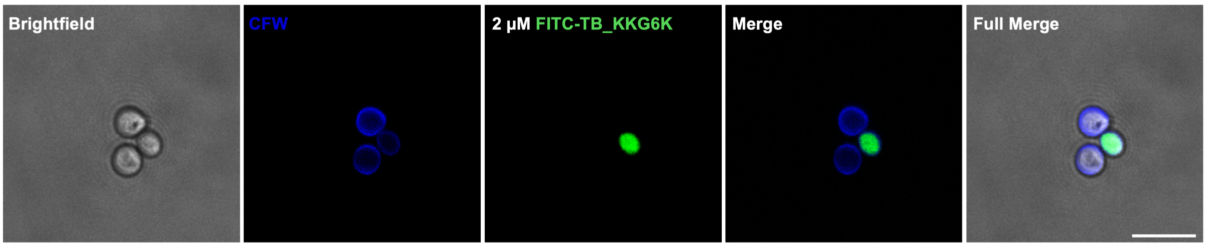


**Figure S2.** Laser scanning microscopic images of *C. albicans* exposed to FITC-labelled TB_KKG6K and stained with the cell wall specific dye calcofluor white (CFW). Cells were exposed to 5 µM of CFW in the presence of 2 µM of FITC-TB_KKG6K for 15 min at 30°C. (A) Overview; scale bar, 20 µm. (B) Detail; scale bar, 10 µm.
